# Characterization and Validation of Compressed Sensing for Time-of-Flight MRI Angiography of the Human Brain at 3T and 7T

**DOI:** 10.64898/2026.04.22.720171

**Authors:** Gustaf Rådman, Xiaole Z. Zhong, Mrinmayi Kulkarni, Valentina Perosa, Jacob J. L. Matthews, Martina F. Callaghan, Emrah Duzel, Dorothea Hammerer, Grazia Daniela Femminella, J. Jean Chen, Rosanna K. Olsen

**Author notes:** **Corresponding author:** Rosanna Kathleen Olsen, J. Jean Chen. These authors contributed equally (Gustaf Rådman, Xiaole Z. Zhong). These authors contributed equally (J. Jean Chen, Rosanna K. Olsen).

## Abstract

**Background:** Cerebral vasculature is a key biomarker of brain health, and time-of-flight (TOF) magnetic resonance angiography (MRA) provides noninvasive assessment of vascular anatomy. However, conventional TOF-MRA requires long scan times, increasing patient burden and susceptibility to motion artifacts. Compressed sensing (CS) offers a feasible acceleration strategy.

**Purpose:** To quantitatively evaluate CS acceleration in TOF-MRA at 3T and 7T using automated whole-FOV vascular segmentation and semi-automatic segmentation of representative vessels.

**Study type:** Prospective

**Population:** 23 healthy human participants (3T) and 8 healthy human participants (7T).

**Field Strength/Sequence:** CS TOF-MRA (CS factors 4 and 8 at 3T; 8 at 7T) was compared against non-accelerated (CS0) TOF-MRA.

**Assessment:** Visual comparison and vascular segmentation were performed using automated whole-FOV methods and semi-automatic segmentation of the posterior cerebral artery and anterior choroidal artery.

**Statistical Tests:** Contrast-to-noise ratio (CNR), voxel count, and vessel diameter were assessed using two-tailed paired t-tests.

**Results:** Whole-FOV CNR differed significantly across CS factors at 3T (CS0 > CS4: p < 0.001, d = 0.77; CS0 < CS8: p = 0.008, d = 0.36; CS4 < CS8: p < 0.001, d = 1.11) and 7T (CS0 < CS8: p = 0.002, d = 0.54), with semi-automatic segmentation yielding consistent findings (p < 0.01 for all comparisons). The diameter measurements for segmented vessels are also higher with high CS-factors (PCA 7T: left: *p* = 0.006, *d* = 0.93, right: *p* = 0.045, *d* = 0.43; AChA 7T: left: *p* < 0.001, *d* = 0.66, right: *p* = 0.009, *d* = 1.06; PCA 3T: *p* < 0.001 for all comparison *d*_Left_ = 0.52 (CS0 vs. CS4), 0.56 (CS4 vs. CS8), 1.11 (CS0 vs. CS8) and *d*_Right_ = 0.78 (CS0 vs. CS4), 0.57 (CS4 vs. CS8), 1.17 (CS0 vs. CS8)).

**Data Conclusion:** CS shows promise for enhancing clinical applicability of TOF-MRA, with advantages most pronounced at 7T.

## 1. Introduction

The cerebral vasculature plays a critical role in supplying nutrients and clearing metabolic waste from the brain (1, 2). Cerebrovascular damage or dysfunction may serve as an early biomarker of subsequent brain function decline and can be predictive of the onset of neurodegenerative diseases (3). Additionally, research has suggested that differences in vascularization patterns themselves can play a decisive role for structural integrity. For example, specific hippocampal vascularization patterns have been shown to be protective against structural and functional decline related to cerebral small vessel disease (4, 5). Furthermore, accurate modeling of susceptibility effects in cerebral macrovasculature is crucial for improving the specificity of functional magnetic resonance imaging (fMRI), thereby enhancing our understanding of brain organization and function, and ultimately advancing its clinical applications (6, 7). Current methods for non-invasive imaging of cerebrovascular anatomy remain limited, despite the clear scientific and clinical importance of accurate, reliable, and efficient approaches.

Time-of-flight (TOF) magnetic resonance angiography (MRA), which leverages the in-flow effect of blood as an endogenous contrast agent, is a widely used noninvasive approach for imaging the anatomy of the cerebral vasculature. Compared with susceptibility-weighted (SW) MRA, TOF-MRA provides superior delineation of the arterial network, particularly for arteries and vessels located near air–tissue boundaries (8, 9). However, as with other MRI techniques, TOF-MRA is fundamentally limited by the trade-off between acquisition time and spatial resolution, as achieving high-resolution imaging requires considerably prolonged scan times (10, 11). In fact, prolonged acquisition time, which is necessary to achieve high resolution, increases susceptibility to motion artifacts, reducing the reliability of vascular measurements (11). Moreover, longer scan times directly translate into higher costs and burden on patients, limiting clinical applicability (12).

Compressed sensing (CS) has been proposed as a strategy to accelerate TOF-MRA acquisitions (13). CS relies on incoherent undersampling of k-space data that is particularly suited to accelerated imaging of spatially sparse objects, in this case blood vessels. This approach has been successfully demonstrated in vessels outside the brain (14), where vascular anatomy is relatively simple and flow rates larger (15, 16). However, the cerebral vasculature is substantially more complex and dense, and flow rates of major brain vessels are lower than those of the torso or leg (17, 18), raising questions about the level of contrast that can be captured using CS.

Some prior studies have evaluated the performance of CS-based TOF-MRA by direct comparison with conventional TOF-MRA in the human brain. Overall, CS-based TOF-MRA, while substantially reducing acquisition time, have shown comparable sensitivity, specificity, and accuracy to conventional approaches at both 3T (13, 19–21) and 7T (22–24). However, there is conflicting evidence for 3T, with image quality highly dependent on the CS acceleration factor and vascular size. Specifically, Ding and colleagues (25) found that arteries across a range of sizes were reliably visible at CS2 and CS4 (where CS2 denotes 2-fold acceleration, CS4 denotes 4-fold acceleration, etc.), that smaller arteries partially lost visibility at CS6, and that only larger arteries remained sufficiently visible at CS8 and CS10.

Although the overall impression is that CS provides important benefits for TOF-MRA in cerebrovascular imaging, previous investigations present some limitations. First, most evaluations have relied on manual segmentations of only major vessels—primarily those in the Circle of Willis, vertebral arteries (VA), and the basilar artery (BA)—limiting generalizability. Moreover, most assessments have been qualitative (20, 21), with only a few incorporating quantitative metrics such as contrast to noise ratios (13, 19, 25) or signal intensity spread functions (23). Complementing previously published evaluations with a more systematic, whole-field-of-view (FOV) evaluation is essential to address current gaps, especially for advancing vascular biomarkers (26, 27) and biophysical brain models (6). Moreover, many previous studies focused on low–spatial-resolution TOF-MRA (13, 19, 25), and their evaluations may not translate to high-resolution imaging, where CS could be even more indispensable given the longer scan times required.

In this study, we combine whole-FOV automated segmentation of cerebral vasculature with semi-automatic segmentation specifically targeting the posterior cerebral artery (PCA) and anterior choroidal artery (AChA) with the aim to comprehensively and quantitatively assessing CS performance in cerebral vasculature at both 3T and 7T. For smaller vessels, we chose to evaluate the PCA and AChA for two reasons. One is that they both provide blood supply to the hippocampus, making them of particular interest to researchers focused on memory and in neuropathologies affecting the hippocampal formation such as Alzheimer’s Disease (4, 28). Second, PCA and AChA differ in their average diameters (29–35), allowing for investigations of size-dependent relationships with CS. With this approach, our findings will promote the broader application of CS-accelerated TOF-MRA and provide a foundation for future clinical and research developments.

## 2. Methods

### 2.1. Participants and Scan Parameters

3T sub-study: A total of 23 participants were imaged at (REDACTED for De-identification purposes) (3T site). After exclusion of two participants with incomplete scans, one participant with severe imaging artifacts, and one participant with incidental findings, 19 participants were included in the 3T dataset (7 males and 12 females, aged 20 to 56). All scans were collected on a Siemens Prisma 3T equipped with a 64-channel head-and-neck receive coil. The TOF-MRA scans included the following compression settings: no compression (CS0), CS-acceleration factor = 4 (CS4), and CS-acceleration factor = 8 (CS8). Additionally, all subjects and CS factors were imaged using both descending and ascending slab orders to minimize bias due to flow direction. However, visibility of vessels proved too low in scans with the ascending order. Results presented here focus on images acquired using the descending order. Finally, a T1-weighted anatomical image (MPRAGE, sagittal, 224 slices, 0.8 mm isotropic resolution, TE = 2.85 ms, TR = 2000 ms, TI = 950 ms, flip angle = 10°) was collected for each participant.

7T sub-study: A total of 8 participants (all females, aged 23 to 40) were imaged at (REDACTED for De-identification purposes). No exclusions were required in the 7T dataset. All 7T scans were collected with Siemens Magnetom Terra equipped with a Nova Medical 8-channel transmit/32-channel receive coil (operates in a quadrature-like “TrueForm”). TOF-MRA was acquired without compression (CS0) and with a CS-acceleration factor of 8. A T1-weighted anatomical image (MPRAGE, sagittal, 224 slices, 0.7 mm isotropic resolution, TE = 3 ms, TR = 2200 ms, TI = 1070 ms, flip angle = 8°) was collected for each participant.

The detailed scan parameters for both 3T and 7T scans are listed in **Table 1**. The FOV for both datasets was set to capture major well defined cerebral arteries (see **Figure 2**). The study was approved by the Research Ethics Boards (REB) at the 3T site (REDACTED for De-identification purposes) (REB 22-32) and at the 7T site (REDACTED for De-identification purposes) (13061/002). All participants provided written informed consent prior to beginning the study. The study was conducted in accordance with the Declaration of Helsinki.

**Table 1.** Detailed scan parameters for 7T and 3T datasets.

| Field Strength | 7T |  | 3T |  |  |
| --- | --- | --- | --- | --- | --- |
| Compression factor | CS0 | CS8 | CS0 | CS4 | CS8 |
| Orientation | Transverse |  | Oblique Transverse |  |  |
| Resolution | 0.28 mm<sup>3</sup> isotropic |  | 0.3 mm<sup>3</sup> isotropic |  |  |
| Slabs and slices | 5 slabs, 48 slices per slab |  | 3 slabs, 56 slices per slab |  |  |
| Flow direction | Foot >> Head |  | Foot >> Head |  |  |
| TR (ms) | 22.0 | 51.9 | 28.7 | 36.3 | 36.3 |
| TE (ms) | 4.59 | 5.61 | 4.62 | 5.08 | 5.08 |
| Flip angle (deg) | 23 | 20 | 18 | 20 | 20 |
| GRAPPA factor | 3 | - | 2 | - | - |
| Scan time (min) | 18:40 | 8:13 | 27:55 | 8:38 | 4:41 |

**Figure 1.**
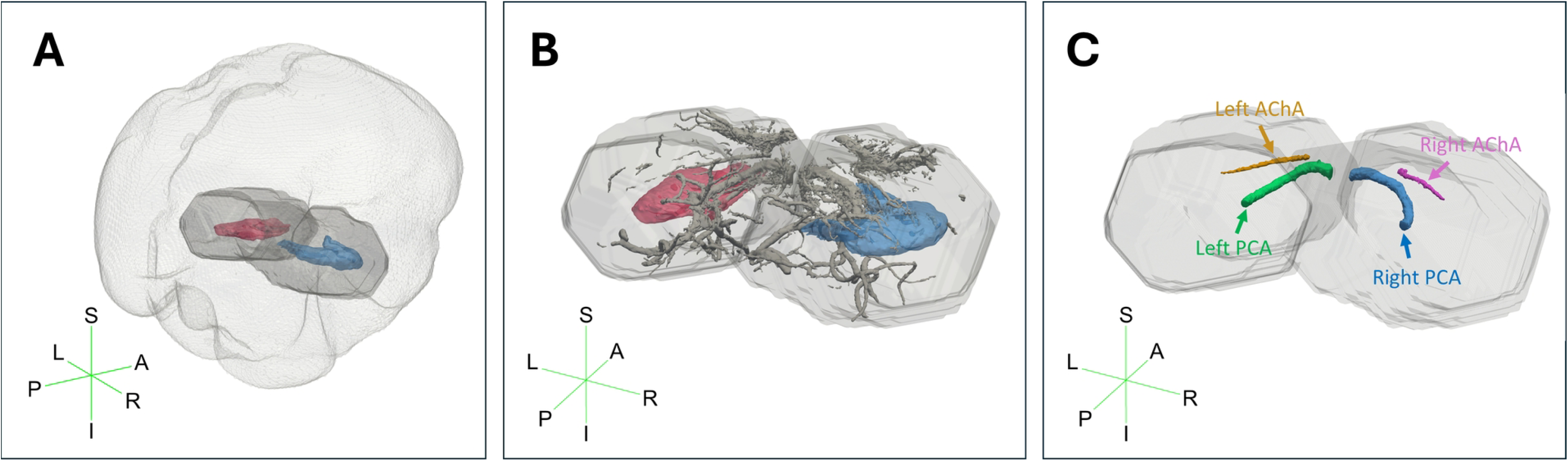
Overview of semi-automatic segmentation of arteries around hippocampal formation at 7T in a representative participant. (A) Regions of interest (ROI) were extracted using dilated masks, seen in darker grey, of left hippocampus (red) and right hippocampus (blue) applied to the TOF MRA. (B) Automated segmentation was applied to extract vasculature in the ROI, seen in grey. (C) Manual labelling was performed to extract the most anterior available segment of left PCA (green), right PCA (blue), left AChA (pink) and right AChA (orange). In 3T, only PCA was segmented due to inconsistent visibility of AChA. Depictions are for illustrative purposes, generated by visualizing mesh surfaces generated for each segmented vessel structure in ParaView (39). Rendered hippocampal masks used for illustrations were made using Automatic Segmentation of Hippocampal Subfields (ASHS; (40)). S=Superior, A=Anterior, R=Right, I=Inferior, P=Posterior, L=Left.

**Figure 2.**
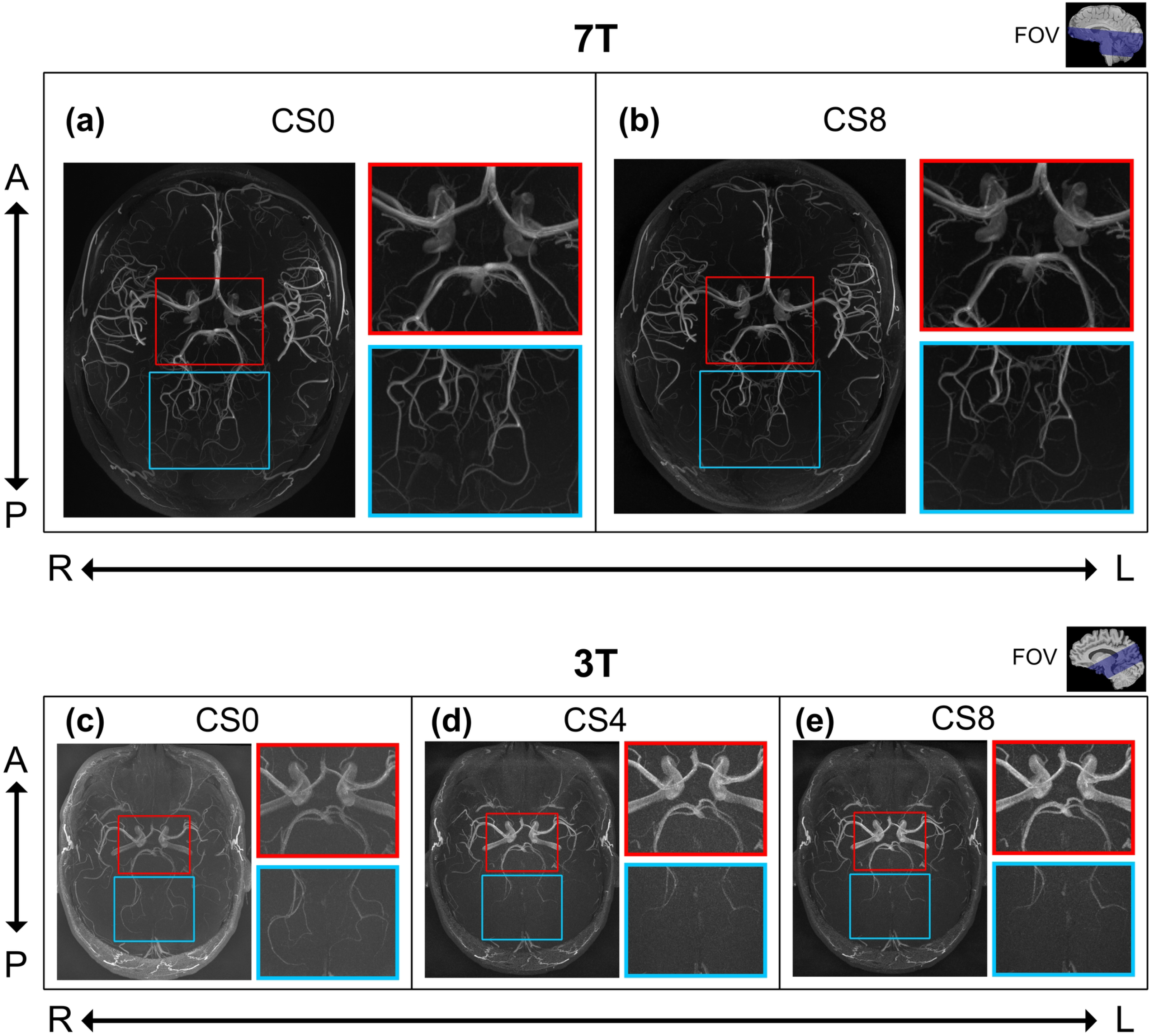
Visual comparison across compression (CS) factors for 7T and 3T. Axial slices of whole-FOV vasculature of 7T and 3T were obtained from representative participants selected from the 7T and 3T datasets, respectively. For 7T MRI, the compression factors included CS0, i.e. uncompressed (a) and CS8 (b), whereas for 3T MRI, the compression factors included CS0 (c), CS4 (d) and CS8 (e). Red boxes provide a zoomed-in visualization of the Circle of Willis. Blue boxes provide a zoomed-in visualization of more distal branches. A = Anterior, P = Posterior, R = Right, L = Left. FOV = Field of View

### 2.2. Image Preprocessing

The preprocessing steps for the 3T and 7T datasets were identical. The anatomical T1-weighted scans as well as the TOF-MRA images were converted to NIFTI format using the *dcm2niix_afni* program in Analysis of Functional NeuroImages (AFNI v21.2.10) (36). All analyses of TOF-MRA scans were conducted in participants’ native space (i.e., without normalization to a template image). T1-weighted scans were segmented using the *recon-all* function in FreeSurfer (v7.1.0; Fischl, 2012), from which hippocampal masks were extracted (from the aseg.mgz segmentation). Additionally, skullstripped versions of the T1-weighted scans were generated using the @SSwarper pipeline in AFNI.

All TOF-MRA scans were aligned to the T1-weighted scan for each subject using functions in Advanced Normalization Tools (ANTs) (37). To prevent the TOF-MRA scans from being downsampled to the resolution of the T1-weighted scans during alignment, the transformation of the TOF-MRA to the T1-weighted scans was first computed using the *antsRegistrationSyNQuick* function. The T1-weighted scans were then upsampled to the resolution of the TOF-MRA images. Finally, the computed transformations were applied to the TOF-MRA scans using the *antsApplyTransforms* function with the upsampled T1-weighted scans as references. For one subject in the 3T dataset co-registration failed to pass visual quality assessment, and manual correction was applied using the manual registration tools in ITK-SNAP. Next, the TOF-MRA scans were skull-stripped. Specifically, binarised masks of the skull-stripped anatomical T1-weighted scans were eroded by 2 mm. These masks were then applied to the aligned TOF-MRA scans using *3dcalc* in AFNI. Finally, the resulting TOF-MRA scans were corrected for bias field (i.e. receive) inhomogeneities using *N4BiasFieldCorrection* in ANTS.

### 2.3. Whole-FOV Automatic Vascular Segmentation

Segmentation of whole-FOV vasculature was automated using a custom pipeline in MATLAB (MATLAB 2021b) (MathWork Inc., Natick, MA, United States). First, a mask was generated through signal intensity thresholding, only retaining voxels above the 98th percentile for 3T and 99th percentile for 7T. Next, connectivity filtering was applied using the *bwconncomp* function from the Image Processing Toolbox. Specifically, segmented structures were only considered vessels if they consisted of 40 or more voxels connected in space.

### 2.4. Semi-automatic Vascular Segmentation of PCA and AChA

To bolster the automated segmentation results, we also applied semi-automated segmentation to a sub-set of vessels. For this we chose the PCA and AChA, two widely studied arteries given their involvement in the hippocampal blood supply system (4, 38). We started by identifying the relevant vascular region using the hippocampal masks derived from FreeSurfer 7 (Fischl, 2012) which were dilated using AFNI (*3dmask*) by an average of 7-10 mm in each direction to encompass the surrounding area. Visual inspection was applied to each mask to ensure they included all available segments of the target vessels. To segment the vessels, we first utilized the automatic pipeline for segmentation described for whole-FOV analysis (see 2.3. Whole-FOV Automatic Vascular Segmentation), with two modifications. First, to further enhance the amount of vasculature covered, we applied a 98th percentile threshold for both 7T and 3T. Additionally, the connectivity filtering threshold was individualized to each scan. For more details of the procedure, please see Supplementary Materials Note 1.

In the resulting segmentations, a tracer (G.R.) familiar with the vascularization of the region manually labelled the portions of the mask belonging to PCA and AChA in the 7T TOF-MRA scans. In the 3T TOF-MRA data, only the PCA was labelled, due to limited visibility of AChA across participants. **Fig. 1** provides an illustration of the segmentation procedure. Because the segmentation was applied in skull-stripped images, the most anterior segments of PCA and AChA were sometimes excluded. Four subjects with 3T scans were excluded because too much of the anterior segment was missing, resulting in a final N = 15. For PCA, the arteries were segmented starting from the most anteromedial segment and ending at its terminal bifurcation leading into the parieto-occipital artery and calcarine artery. For AChA, vessels were segmented starting from the most anterior available slice and moving as far as possible posteriorly. Crucially, across both vessels, special care was taken to consistently start and end the segmented area on the same slice for all scans (e.g., CS0, CS8) of a given participant.

The final arterial masks were subjected to a procedure to derive diameter estimates for each point along the vessel, allowing us to compare obtained diameter estimates to previously reported vessel sizes. The extraction of diameters was performed using functions from the Image Processing Toolbox in MATLAB. Specifically, segmentations were first skeletonized using *bwskel* to extract centerlines, followed by detection of vessel boundaries using *bwmorph3* (with remove function). Finally, calculations of the vessel radius were performed based on the average distance between each centerline voxel and the nearest vascular boundary using *bwdist*. This value was doubled to get the diameter, and the median diameter over all centerline voxels was used as a summary diameter value for the vessel being analyzed. For 3T, morphological closing was applied before extractions of diameters due to occasional intra-vessel signal loss which complicated reliable diameter extraction (for more details, see Supplementary Materials Note 2 and Supplementary **Fig. S1-S2**).

### 2.5. Metrics

#### 2.5.1. Whole-FOV Vasculature

For the quantitative comparison, two metrics representing different aspects of vasculature were defined for the segmented vasculature. The metrics were:

● Count of vascular voxels: An indicator of sensitivity to the specific vessel.
● Contrast to noise ratio (CNR; Eq. 1): An indicator of the contrast induced by the TOF effect, which is commonly used to assess image quality (25). CNR values were min-max normalized (range 0-1) collapsed across all CS factors.

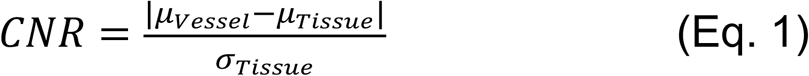

μ_Vessel_ is the mean signal intensity of the vessel, μ_Tissue_ is the mean signal intensity of the brain tissue and **σ**_Tissue_ is the standard deviation of the signal intensity of the brain tissue.

Two-tailed paired t-tests were used to determine whether there were significant differences between compression factors (*α* = 0.05). Statistical analyses were conducted in R 4.5.2 (41). The statistical analysis of 3T and 7T scans was conducted separately. Effect sizes were also calculated using Cohen’s *d* (42), where a *d* higher than 0.2 is considered a small effect, a *d* higher than 0.5 is considered a medium effect, and a *d* higher than 0.8 is considered a large effect (43).

#### 2.5.2. PCA and AChA segmentations

For PCA (3T and 7T) and AChA (7T), effects of CS were evaluated using CNR (as defined earlier) and median diameter values separately for each vessel and hemisphere. Two-tailed paired t-tests (*α* = 0.05), conducted in R, were used to assess differences between compression factors, with 3T and 7T data analyzed separately. Effect sizes were calculated using Cohen’s *d* (*42*).

## 3. Results

### 3.1. Visual Comparison of Whole-FOV Vascular Sensitivity

Whole-FOV vasculature from representative participants (one for 7T and one for 3T) are shown in **Fig. 2**. In the case of 7T, CS8 and CS0 results are visually comparable, with no clear differences observed either in larger vessels such as those of Circle of Willis (see red boxes, 7T, **Fig. 2**) or in more distal, smaller branches (see blue boxes, 7T, **Fig. 2**). In 3T, larger vessels such as those of Circle of Willis (red boxes, 3T, **Fig. 2**), had comparable visibility across CS factors. However, visibility of smaller, more distal, branches seem slightly better in CS0 scans compared to CS scans (blue boxes, 3T, **Fig. 2**).

### 3.2. Quantitative Comparison of Metrics Extracted from Whole-FOV Segmented Vasculature

The mean and standard deviation for each evaluated metric is presented in **Table 2**. Automatically segmented vasculature was first compared at a whole-FOV level across the acquisition types. **Fig. 3a and 3b** show the vascular voxel count for 7T and 3T data, respectively, whereas **Fig. 3c and 3d** show the contrast to noise ratios for 7T and 3T, respectively. At 7T, the number of voxels identified as vessels was significantly greater for CS8 than CS0 (*p* = 0.002, *d* = 0.54). CS8 also showed a significantly higher CNR than CS0 (*p* < 0.001, *d* = 1.51).

**Figure 3.**
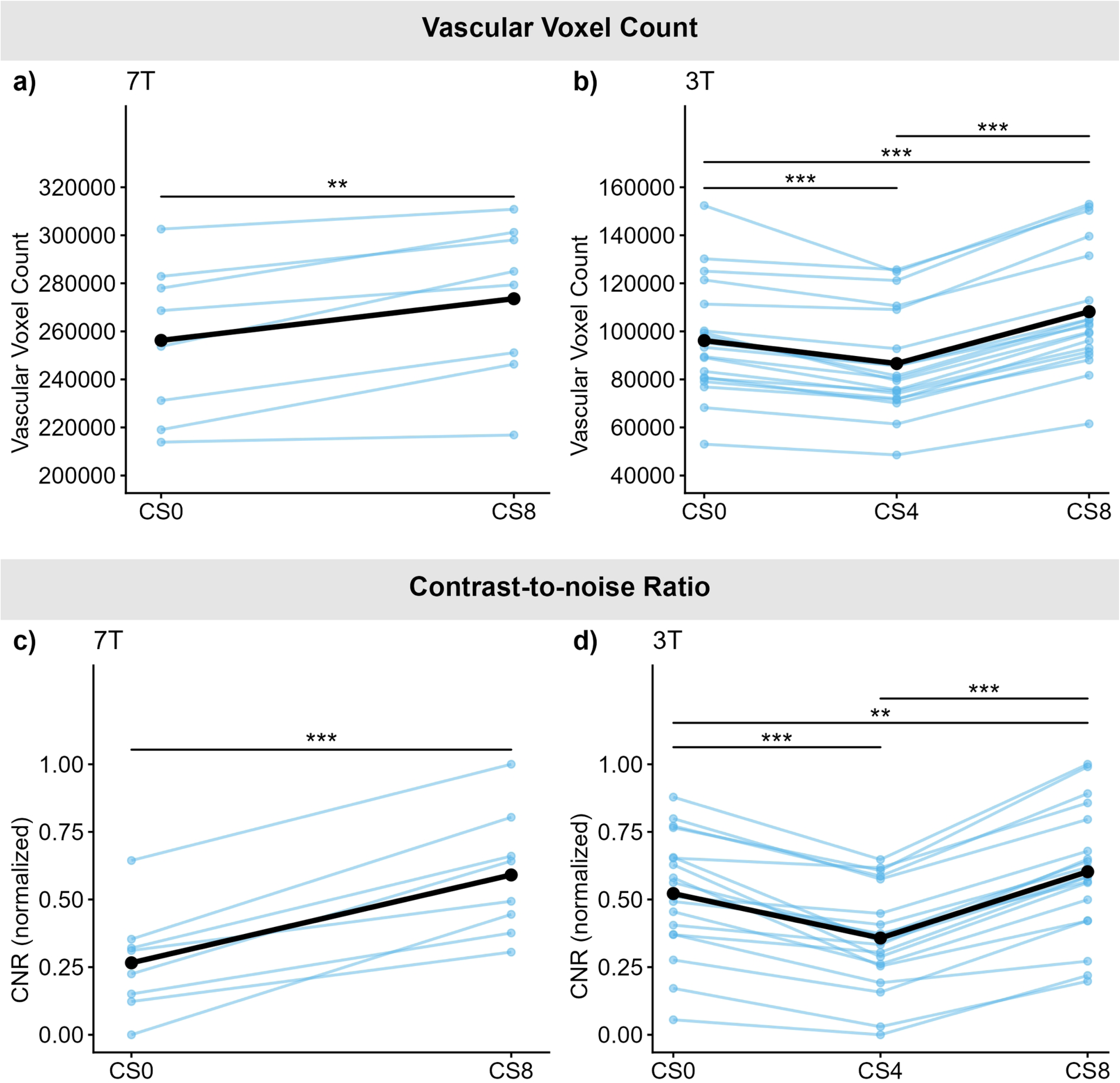
The vascular-voxel counts (a-b) and contrast to noise ratios (CNR) (c-d) for 3T and 7T using whole-FOV automated segmentation. *Left:* 7T; *Right:* 3T. Blue lines represent individual subjects, black lines the average across subjects. Asterisk signs significance assessed using paired t-tests (*: *p* < 0.05; **: *p* < 0.01; ***: *p* < 0.001).

**Table 2.** Mean and standard deviation for all evaluated metrics in whole-FOV analysis and analysis of PCA/AChA by hemisphere. SD: inter-participant standard deviation.

| Field Strength | 7T |  | 3T |  |  |
| --- | --- | --- | --- | --- | --- |
| Compression factor | CS0 | CS8 | CS0 | CS4 | CS8 |
| *Whole-FOV* |  |  |  |  |  |
| Mean CNR ± SD | 0.27 ± 0.19 | 0.59 ± 0.23 | 0.52 ± 0.22 | 0.36 ± 0.19 | 0.60 ± 0.24 |
| Mean Voxel-count ± SD ($\times 10^5$) | 2.56 ± 0.32 | 2.74 ± 0.33 | 0.96 ± 0.24 | 0.87 ± 0.22 | 1.08 ± 0.26 |
| *PCA Left* |  |  |  |  |  |
| Mean CNR ± SD | 0.34 ± 0.16 | 0.52 ± 0.13 | 0.30 ± 0.14 | 0.59 ± 0.18 | 0.77 ± 0.19 |
| Mean diameter ± SD | 2.03 ± 0.12 | 2.12 ± 0.07 | 1.84 ± 0.17 | 1.93 ± 0.20 | 2.04 ± 0.20 |
| *PCA Right* |  |  |  |  |  |
| Mean CNR ± SD | 0.50 ± 0.12 | 0.75 ± 0.15 | 0.30 ± 0.11 | 0.55 ± 0.11 | 0.76 ± 0.09 |
| Mean diameter ± SD | 2.10 ± 0.17 | 2.17 ± 0.17 | 1.83 ± 0.14 | 1.94 ± 0.15 | 2.03 ± 0.15 |
| *AChA Left* |  |  |  |  |  |
| Mean CNR ± SD | 0.32 ± 0.10 | 0.57 ± 0.21 | - | - | - |
| Mean diameter ± SD | 0.97 ± 0.08 | 1.02 ± 0.08 | - | - | - |
| *AChA Right* |  |  |  |  |  |
| Mean CNR ± SD | 0.34 ± 0.16 | 0.64 ± 0.31 | - | - | - |
| Mean diameter ± SD | 0.98 ± 0.09 | 1.07 ± 0.07 | - | - | - |

At 3T, differences in vascular-voxel count across CS factors were statistically significant for all three comparisons (CS0 vs CS4: *p* < 0.001, *d* = 0.42; CS0 vs CS8: *p* < 0.001, *d* = 0.49; CS4 vs CS8: *p* < 0.001, *d* = 0.91), with CS8 showing the highest vascular-voxel count, followed by CS0, and CS4 the lowest. CNR values also demonstrated statistically significant differences across CS factors (CS0 vs CS4: *p* < 0.001, *d* = 0.77; CS0 vs CS8: *p* = 0.008, *d* = 0.36; CS4 vs CS8: *p* < 0.001, *d* = 1.11), with CS8 having the highest CNR, followed by CS0, and CS4 once again the lowest. See **Fig. S3** for the numerator (vascular-tissue contrast) and denominator (noise as standard deviation of tissue signal) of the CNR calculation plotted separately.

### 3.3. Quantitative Comparison of Metrics Extracted from PCA and AChA Segmentations

#### 3.3.1 PCA and AChA in 7T TOF-MRA

Focusing on 7T, **Fig. 4a-d** displays the median diameter values of PCA and AChA vessel measurements across CS factors and **Fig. 4e-h** the CNR values across CS factors.

**Figure 4.**
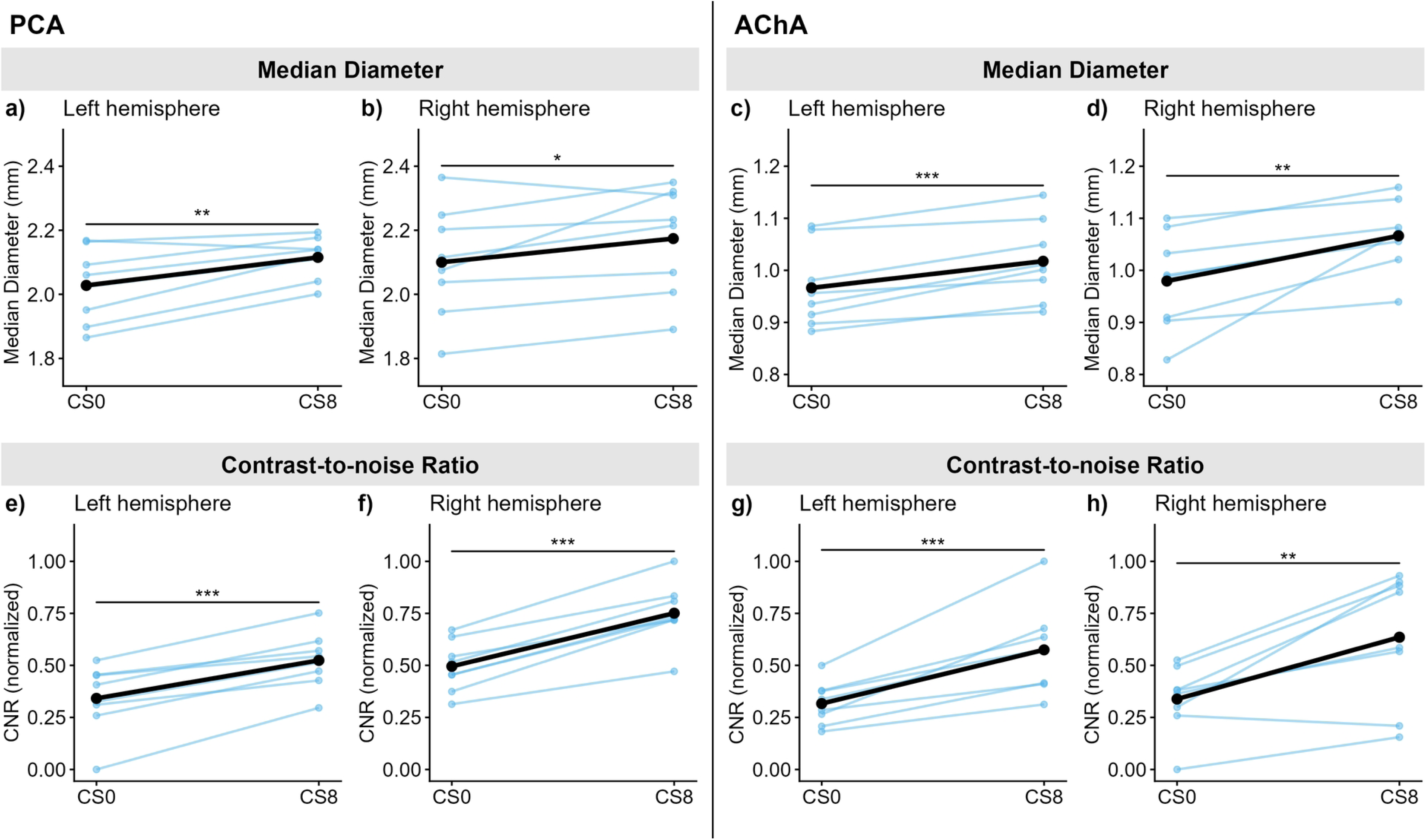
Median diameter (a-d) and contrast to noise ratios (CNR) (e-h) for PCA and AChA at 7T. *Left*: Posterior Cerebral Artery (PCA), *Right*: Anterior Choroidal Artery (AChA). Blue lines represent individual participants, black lines the average across subjects. Asterisk signs significance assessed using paired t-tests (*: *p* < 0.05; **: *p* < 0.01; ***: *p* < 0.001).

For the PCA, there was a statistically significant difference in the median diameter comparing CS0 and CS8 (left: *p* = 0.006, *d* = 0.93; right: *p* = 0.045, *d* = 0.43), with CS8 having higher median diameters than CS0. This was mirrored in CNR values, which were significantly higher for CS8 than CS0 (left: *p* < 0.001, *d* = 1.23; right: *p* < 0.001, *d* = 1.84).

Results for AChA show a similar pattern. There was a statistically significant difference in the median diameters associated with CS0 and CS8 (left: *p* < 0.001, *d* = 0.66; right: *p* = 0.009, *d* = 1.06), with higher median diameters for CS8 than CS0. Likewise, there was a statistically significant difference in CNR values comparing CS8 and CS0 in both hemispheres (left: *p* < 0.001, *d* = 1.52; right: *p* = 0.004, *d* = 1.22), with higher CNR values for CS8 compared to CS0. See **Fig. S4** for the numerator (vascular-tissue contrast) and denominator (noise as standard deviation of tissue signal) of the CNR calculation plotted separately.

#### 3.3.2. PCA in 3T TOF-MRA

Given the lack of consistent visibility of AChA in 3T TOF-MRA, only PCA results were compared across compression factors at 3T. **Fig. 5a-b** show the median diameter values and **Fig. 5c-d** the CNR values across CS factors. We found in both hemispheres that median diameters were highest for CS8, followed by CS4, and lowest for CS0. All comparisons were statistically significant at *p* < 0.001 with effect sizes *d*_Left_ = 0.52 (CS0 vs. CS4), 0.56 (CS4 vs. CS8), 1.11 (CS0 vs. CS8) and *d*_Right_ = 0.78 (CS0 vs. CS4), 0.57 (CS4 vs. CS8), 1.17 (CS0 vs. CS8).

**Figure 5.**
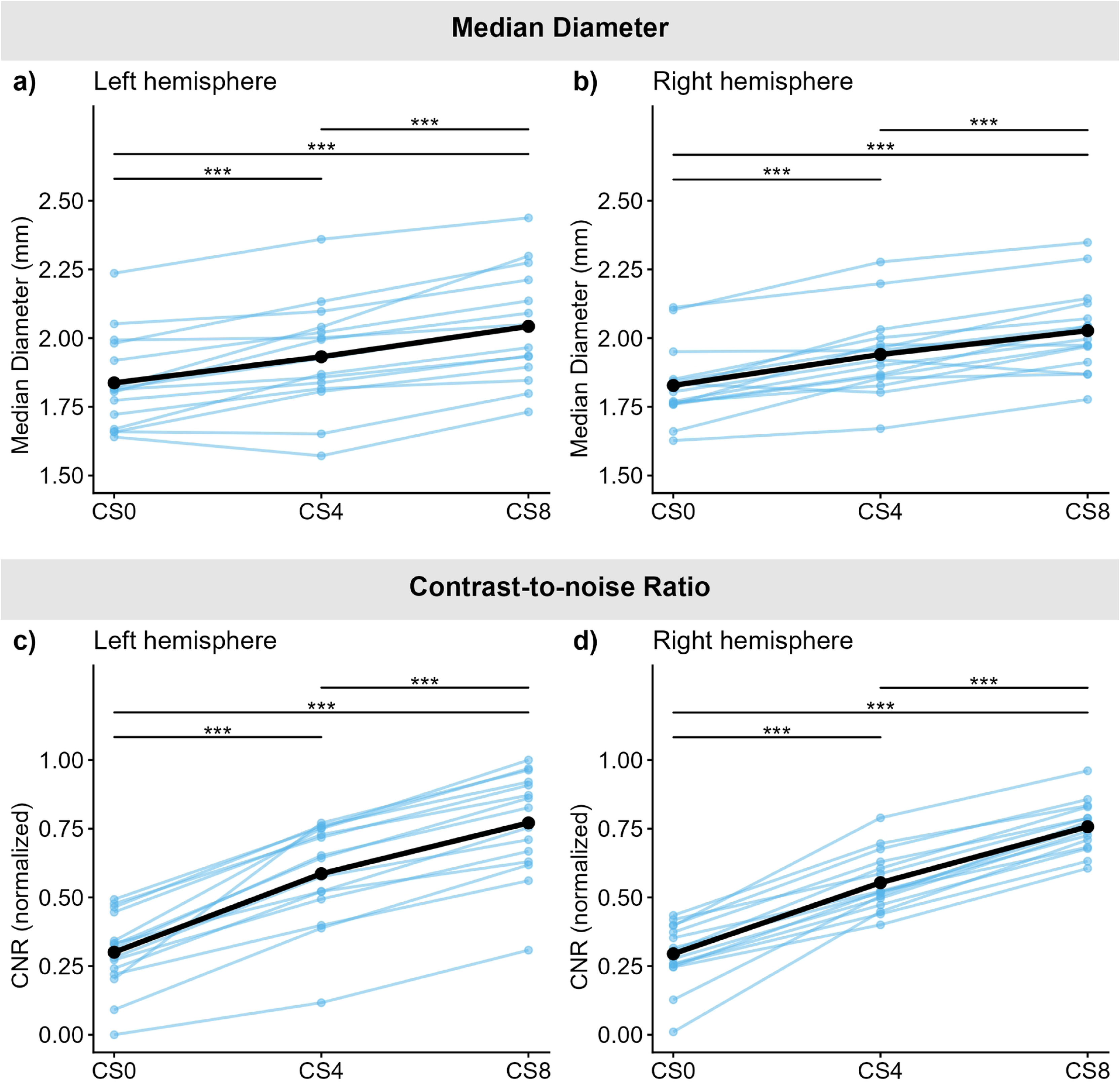
Median diameters (a-b) and contrast to noise ratios (CNR) (c-d) for PCA at 3T. Blue lines represent individual participants, black lines the average across subjects. Asterisk signs significance assessed using paired t-tests (*: *p* < 0.05; **: *p* < 0.01; ***: *p* < 0.001)

The same pattern emerged for CNR values, with CS8 having the highest CNR values, followed by CS4, and CS0 the lowest. All comparisons were statistically significant at *p* < 0.001 with effect sizes *d*_Left_ = 1.79 (CS0 vs. CS4), 0.97 (CS4 vs. CS8), 2.82 (CS0 vs. CS8) and *d*_Right_ = 2.27 (CS0 vs. CS4), 2.09 (CS4 vs. CS8), 4.58 (CS0 vs. CS8). See **Fig. S5** for the numerator (vascular-tissue contrast) and denominator (noise as standard deviation of tissue signal) of the CNR calculation plotted separately.

## 4. Discussion

CS can play an important role in accelerating MRA acquisition, enhancing its feasibility for both clinical and research applications by drastically reducing scan times. In this study, we aimed to evaluate the effects of CS on image quality in 7T and 3T TOF-MRA at a whole-FOV level and within two selected vessels of clinical and research relevance – the posterior cerebral artery (PCA) and anterior choroidal artery (AChA). Our findings suggest an undeniable benefit of CS at 7T, reducing scan times while maintaining image quality. Conversely, CS applied to 3T TOF-MRA resulted in improved performance using a compression factor 8 but inconsistent effects with a compression factor 4. Together, these findings suggest that the performance of CS TOF-MRA compared to CS0 TOF-MRA depends on both CS factor and field strength.

### 4.1. Effect of Compressed Sensing on 7T TOF-MRA

CS was clearly beneficial at 7T. Qualitatively, CS TOF-MRA at 7T produced highly comparable images to that of CS0 (CS0) imaging (**Fig. 2**), consistent with previous studies (22–24). Quantitatively, CS8 scans displayed a consistently higher CNR and voxel count in whole-FOV automated segmentations, in keeping with the qualitative evaluation. These findings were corroborated by observing a similar pattern of results measuring CNR and median diameters for PCA and AChA using a semi-automated approach.

By focusing on a set of defined vessels with previously reported diameter values, we were able to assess the accuracy of our diameter measurements across acquisition types. For the PCA, the current study found diameters averaging 2.06 mm in CS0 and 2.14 mm in CS8. These values are below previous estimates derived from microsurgical data averaging around 2.5 mm (31, 32, 34) but higher than many previous studies using angiography. For instance, using CT angiogram, Kocak et al. ((44) found diameters for PCA averaging 1.76 mm. Huck et al. (2025), using 1.5T MRA, reported an average diameter of 1.96 mm for PCA when manually measured by a trained vascular neurologist, but a diameter of 2.46 mm using Express IntraCranial Arteries Breakdown (eICAB), a recently developed toolbox for automated segmentation of the Circle of Willis. Notably, most previous estimates were obtained from the P1 segment, whereas in the current study, PCA diameter was estimated over the full length of its main trunk. Given that there is a tapering effect over the course of segments P1-P3 (34), an analysis of only P1 would likely have resulted in a higher diameter estimate in the current study.

For AChA, diameters in the current study averaged 0.97 mm in CS0 and 1.04 mm in CS8. These values are slightly higher than microsurgical reports at approximately 0.9 mm (29, 31). Moreover, they are higher than previous reports using DSA and CT angiograms which have found diameter values around 0.8-0.9 mm (e.g., (30, 33). To the best of our knowledge, no previous studies have reported diameter values of AChA derived specifically using MRA imaging. The increased estimate relative to microsurgical values can likely be ascribed in part to partial voluming effects slightly inflating diameter estimates, as well as inherent differences in modality. However, compared to previous DSA/CT studies, the current study achieved higher image resolution, suggesting partial voluming cannot account for the observed differences. Rather, we deem it likely that the current study simply managed to capture more vasculature.

Overall, our results suggest that at 7T, CS imaging represents an improvement over CS0 imaging. This could be due to the reduced scan time with CS8, which can minimize motion-related blurring, allowing more vascular voxels to surpass the threshold of detection. The drastically reduced scan time itself (from 18:40 min at CS0 to 8:13 min at CS8) is a major advantage for patients and research participants. Differences in CNR and diameter values may also arise from variations in the point spread function (PSF) comparing CS0 and CS8 (45, 46), with CS8 typically exhibiting a lower peak but wider full-width-half-maximum (FWHM) PSF than CS0 (47). However, the effect of CS on PSF previously reported is small, thus this effect is unlikely to be the primary contributor (47). Rather, in line with a growing number of studies, we suggest that CS is an attractive option for 7T TOF MRA imaging, providing at least comparable imaging at a reduced cost and burden put on participants. Given that in this study, only a single compression factor was evaluated at 7T, future work is encouraged to identify whether a higher or lower compression factor might provide additional benefits, either in terms of improved vascular imaging or reduced scan times.

### 4.2. Effect of Compressed Sensing on 3T TOF-MRA

At 3T, qualitatively, uncompressed scans show a clear benefit in the ability to visualize smaller vessels, with largely comparable visibility of distal branches between CS4 and CS8 (see Blue boxes in **Fig. 2**). Interestingly, unlike previous work (with the same CS factors) showing a monotonic trend with higher CS factors reducing vessel visibility and CNR (25), our quantitative analysis at a whole-FOV level reveals a bitonic dependency on CS factor, where vascular voxel counts and CNR followed a V-shaped trajectory with CS8 outperforming both CS0 and CS4 and CS4 performing least well. In the study by Ding et al. (2021) higher compression factors had a negative impact on CNR, whereas the current study instead showed an initial reduction in CNR followed by an increase when the highest compression factor of 8 was used. In fact, in analysis restricted to the PCA, we observed an increase in CNR values as the CS factor increased from 0 to 8. Similarly, the extracted vascular diameters for the PCA were also greatest for CS8 (2.04±0.20 mm), followed by CS4 (1.94±0.15 mm), with the smallest being for CS0 (1.83±0.14 mm).

A possible explanation for the bitonic behaviour as a function of the CS factor at 3T (whole-FOV analysis) may lie in the trade-off between potential blurring from compressed sensing from reconstruction and from motion artifacts. Ding et al., (2021) used larger voxel sizes (acquisition resolution: 0.52 x 0.8 x 1.35 mm_3_) and shorter scan times (unaccelerated = 7:18; SENSE factor 2 = 3:38; CS4 = 1:57; CS8 = 1.00), making acquisitions less prone to motion artifacts but more prone to the effects of CS-related undersampling. By contrast, the present study used smaller voxels and overall longer scan durations, reversing this trade-off. That is, low CS factors may result in reduced contrast for small vessels through motion blurring, whereas further increasing the CS factor may help restore contrast by mitigating motion artifacts at the expense of introducing CS-related PSF effect. Additionally, it should be noted that the CS0 imaging of the current study, but not CS4 or CS8, incorporated parallel imaging using GRAPPA acceleration, in contrast to the non-accelerated and SENSE-accelerated CS0 configurations employed in previous studies (e.g., (25). This may lead to subtle differences in the noise characteristics compared to non-GRAPPA CS4 and CS8 images.

We observed qualitative benefits of CS0 imaging for detection of more distal branches at 3T, but a clear benefit of CS imaging for larger vessels such as the PCA. This pattern of results may be explained by the sensitivity of compressed sensing to vascular size. Because compressed sensing oversamples low-frequency k-space (48), it is inherently more sensitive to larger vessels than to smaller ones, which agrees with previous reports (25). The PCA, being a relatively large vessel, occupies comparatively lower spatial frequencies and therefore benefits more from CS compared to smaller (higher spatial frequency) vessels. Overall, our findings suggest that CS acquisitions, compared with CS0, are able to better preserve the morphological features of vasculature the size of PCA at 3T, but that achieving comparable imaging of smaller, more distal branches via CS imaging may require further sequence optimization. Of note, there was no clear difference in the visibility of AChA across CS and CS0 scans at 3T, both image types having limited visibility. Similar to 7T, other factors may also play a role in the observed differences of the current study, such as differences in PSF for CS and CS0 scans (47).

### 4.3. Comparison of Compressed Sensing TOF-MRA at 3T and 7T

At 7T, the stronger field facilitates stronger inflow contrast, enabling higher vascular contrast compared to 3T MRI at the same spatial resolution. Although the FOV differed substantially between 7T and 3T acquisitions in the current study (7T: R-L: 175 mm, A-P: 200 mm, F-H: 54 mm; 3T: R-L: 173 mm, A-P: 200 mm, F-H: 44 mm), it is clear by visual inspection that 7T TOF-MRA overall provides much better visibility for smaller vessels (**Fig. 2**). While both 3T and 7T manage to capture cerebral arteries regardless of acquisition type, only 7T manages to consistently capture smaller branches such as the AChA or posterior communicating artery. The intrinsic differences in inflow contrast and motion sensitivity between 3T and 7T also influence their response to compressed sensing. Theoretically, 7T—with a stronger inflow effect and shorter scan time—should benefit less from CS than 3T. Comparing the results for CS0 and CS8 across 7T and 3T does not align with this notion. In terms of voxel count, the effect of CS8 was comparable between 7T and 3T. However, in terms of contrast to noise ratios, CS8 had a larger effect at 7T than at 3T. Furthermore, results isolated to PCA corroborated these findings, demonstrating a larger effect for both CNR and diameter values at 7T compared to 3T. The stronger inflow effect at 7T produces sharper vessel boundaries (approaching a square intensity profile), which contain more low-frequency components than the blurred boundaries at 3T (49, 50), which may contribute to the observed differences.

Although the precise mechanisms remain unclear, these findings imply that the impact of compressed sensing might not only depend on vessel size but also on signal intensity (in-flow effect) and its sharpness, which together determine its spatial frequency representation. Overall, our whole-FOV analysis indicates that compressed sensing improves TOF-MRA at 7T with very little down-side, supporting its further implementation. At 3T, conversely, compressed sensing remains limited in revealing small sized vessels already not visible in CS0 scans but seems to provide benefits for the segmentation of vessels at the size of cerebral arteries. Future studies should consider varying parameters such as voxel size (and associated scan time) and CS factors to better characterize the operational boundary, especially in future inquiry into the use of TOF-MRA at 3T.

### 4.4. Limitations and future directions

This study was performed on a small sample, limiting generalizability. Estimated effect sizes are likely prone to relatively large error margins. However, most effects found were large and consistent in their directionality. Moreover, we only tested a limited set of CS factors and within the scans acquired kept other parameters as constant as possible. Several key questions remain regarding the influence of different factors in determining image quality and detectability of vessels across different sizes, especially for future use of 3T imaging. Specifically, the interaction between voxel size, CS factor and vessel size requires further investigation. Moreover, comparison of our results to previous studies is not fully straightforward due to key differences in acquisition parameters such as voxel sizes and acquisition times (e.g., (25). Additionally, the automatic segmentation pipeline used in this study serves as a simplified approach compared to other tools available that were not suited to apply in the current study (e.g., BrainCharter, which requires whole-brain imaging; (15).

## Conclusion

CS enables faster TOF-MRA acquisitions, reducing both cost and patient burden, thereby increasing clinical applicability. Despite its growing importance, the quantitative validation of CS across different CS factors and whole-FOV vasculature has been limited. In this study, we evaluated the effectiveness of CS using both automated whole-FOV segmentation and semi-automatic segmentation of representative vessels (PCA and AChA) at 3T and 7T. In addition to assessing CNR as a measure of TOF-MRA sensitivity, we compared extracted counts of vascular voxels and compared vascular diameters of select vessels with established reference values. Our results provide quantitative evidence supporting the reliability of CS in preserving vascular morphology, shortening scan times, and largely preserving image quality. This advantage is especially pronounced at 7T. Our results highlight the potential of CS to advance the clinical utility of TOF-MRA.

## Data availability

The code will be made available upon request. The data cannot be shared publicly due to ethics approval restrictions. Data and code can be accessed by contacting the corresponding authors.

## Supplementary Material

### Note 1. Dynamic identification of threshold for connected voxels in semi-automatic segmentations of hippocampal supplying vessels

To further increase our chances of capturing as much vasculature as possible we lowered the intensity threshold used to the 98% percentile for both 3T and 7T. Further, we dynamically adapted the minimum required number of connected voxels used for connectivity filtering. This was done for each scan independently by plotting the distribution of cluster sizes (i.e. number of voxels) of all connected components remaining after intensity thresholding and identifying the kneepoint in this distribution. Next, the value at this kneepoint was used as the minimum required number of connected voxels. To avoid local minima in the distribution, we set the lower bound for this kneepoint to 20 voxels, informed by visual inspection of the histogram distributions over different scans and by visual inspection of initial segmentations attempts.

### Note 2. Sporadic signal loss in 3T TOF-MRA

In 3T TOF-MRA, scans exhibited signal loss frequently in the center of vessels. Example slices with the issue are presented below in **Figure S1** (top and middle row) for the posterior cerebral artery (red voxels). For semi-automatic segmentation of PCA, since the vessel was isolated using manual labelling of the vessel, it was possible here to apply morphological closing of the vessels. Specifically, the PCA label was first dilated by two voxels, followed by erosion by the same amount of voxels, results of which are seen in **Figure S1** (bottom row). There was a significant difference in the amount of morphological closing applied across the three acquisition types (CS0, CS4, CS8) (see **Figure S2**). Specifically, the most morphological closing was applied in CS0, followed by CS4 and CS8.

## Glossary

AChA: Anterior choroidal artery
AFNI: Advanced Functional Neuroimaging toolbox
ANTs: Advanced Normalization Tools
ASHS: Automatic Segmentation of Hippocampal Subfields
CNR: Contrast ratio
CS: Compressed sensing
FOV: Field of view
TOF: Time-of-flight
MPRAGE: Magnetization-prepared rapid gradient echo
MRA: Magnetic resonance angiography
MRI: Magnetic resonance imaging
PCA: Posterior cerebral artery
SWI: Susceptibility weighted imaging
TE: Echo time
TI: Inversion time
TR: Repetition time

---

**Figure S1.**
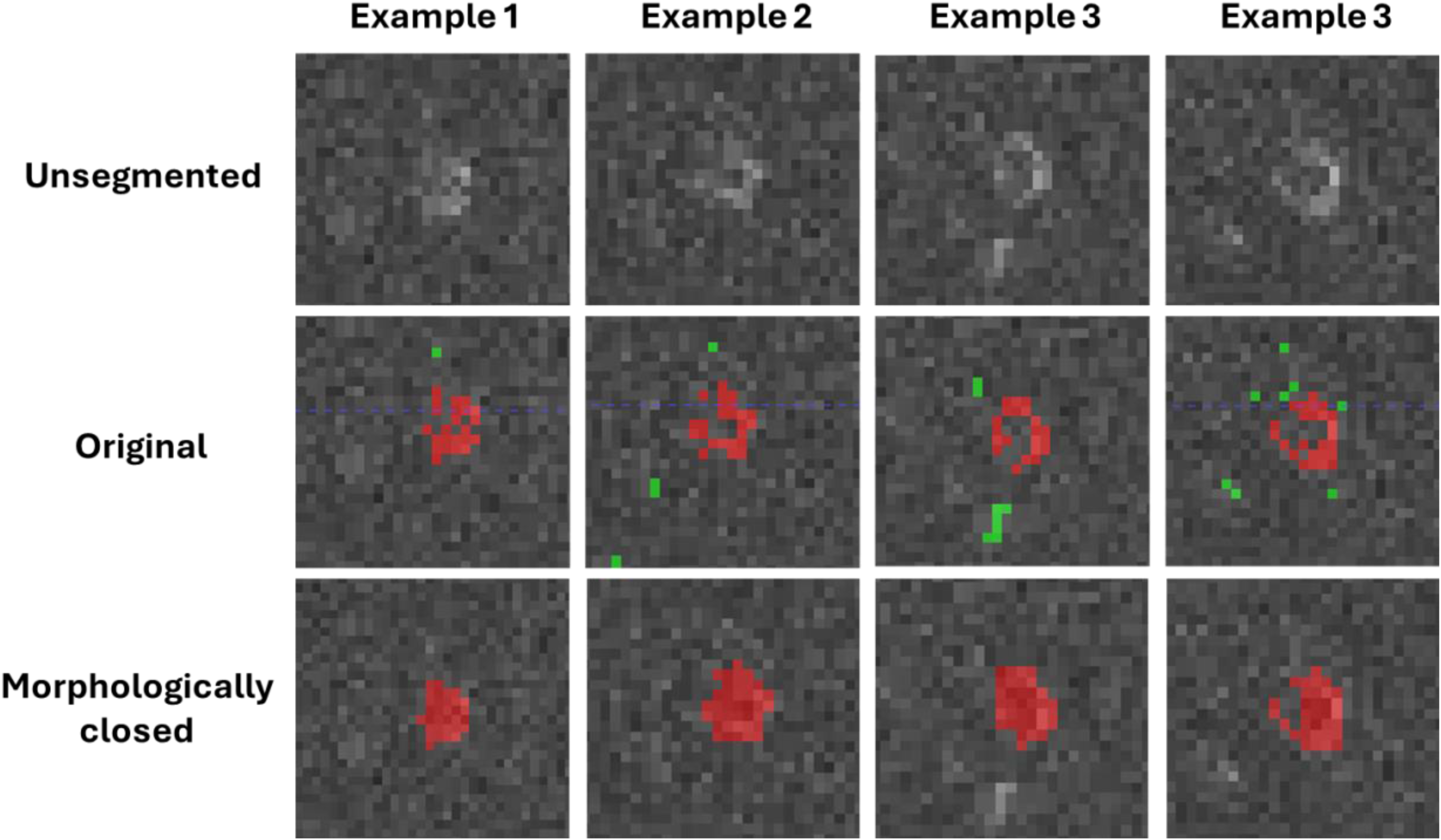
Examples of slices with sporadic signal loss in 3T for the posterior cerebral artery (PCA). Red = PCA. Green = voxels initially included in the automatic vascular segmentation but manually excluded from final vessel segmentation. Morphological closing using dilation followed by erosion was applied to the assisted manual segmentation of PCA (bottom panel)

**Figure S2.**
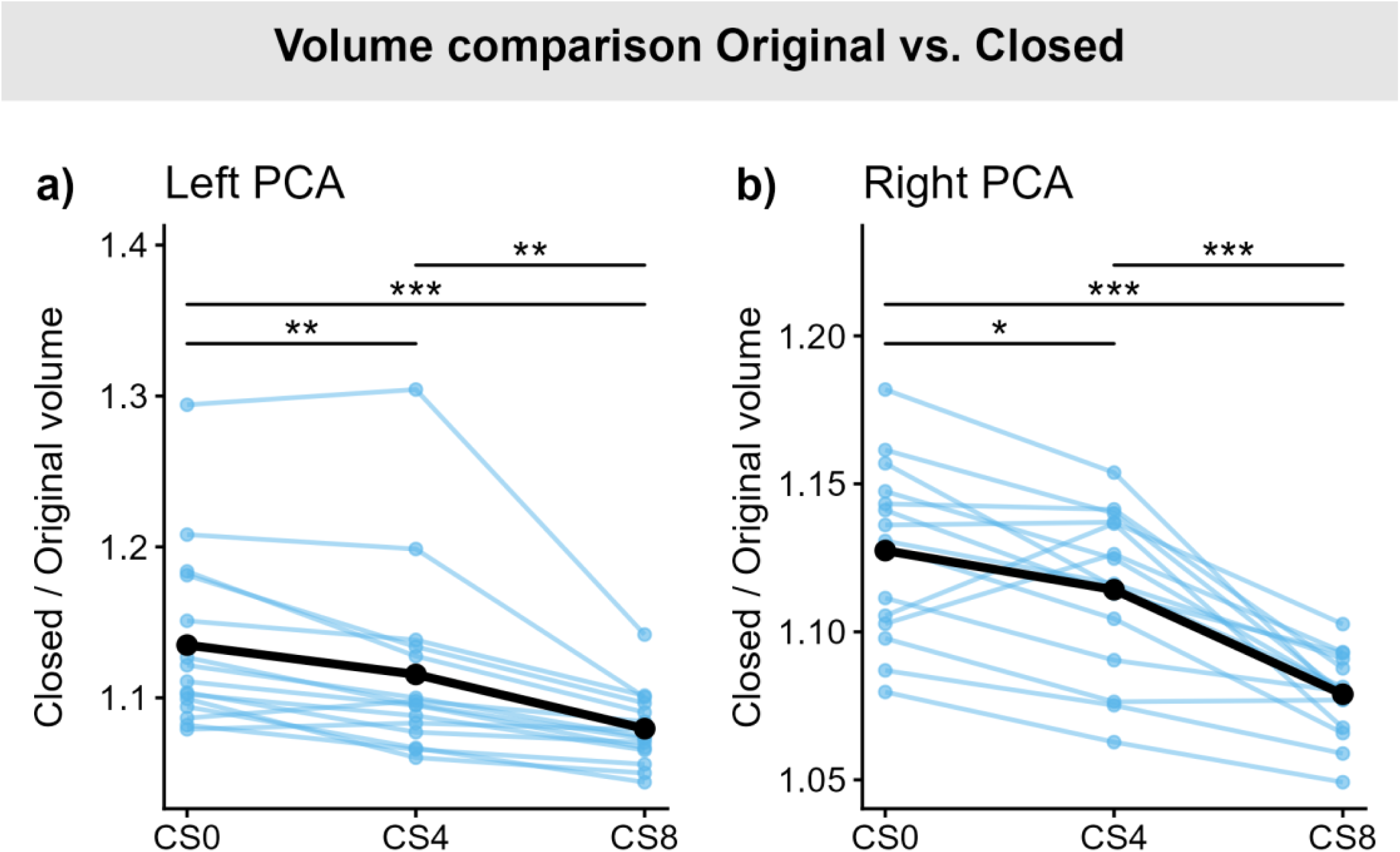
Volume comparison Original versus morphologically closed segmentations of PCA using assisted manual segmentation. Morphological closing added the most voxels for CS0, more voxels for CS4 and the most voxels for CS8. Asterisk signs significance assessed using paired t-tests (*: *p* < 0.05; **: *p* < 0.01; ***: *p* < 0.001).

**Figure S3.**
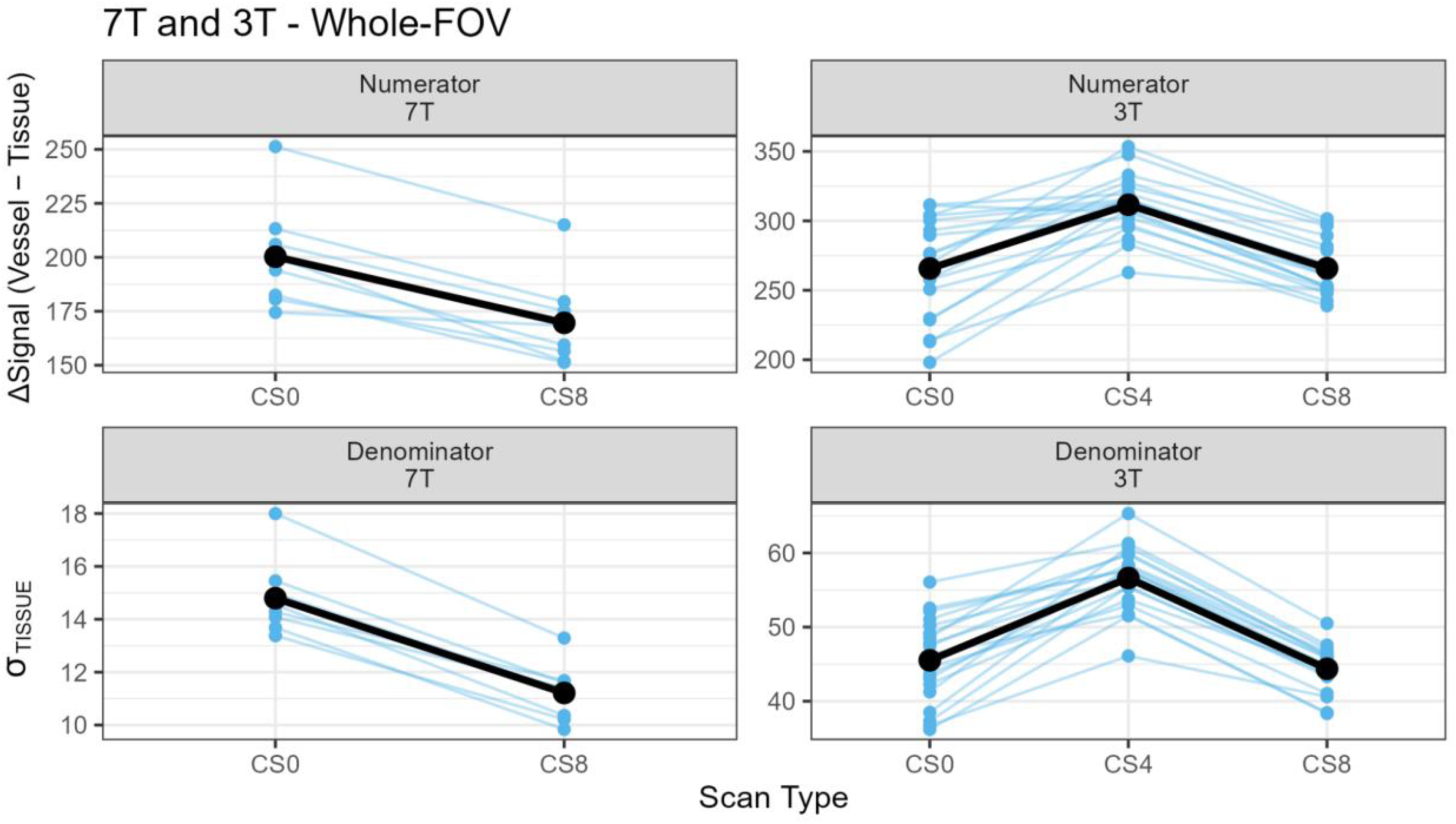
Comparison of acquisition types (CS0 and CS8) for whole-FOV at 7T and 3T separating the effect in numerator and denominator of the contrast-to-noise formula.

**Figure S4.**
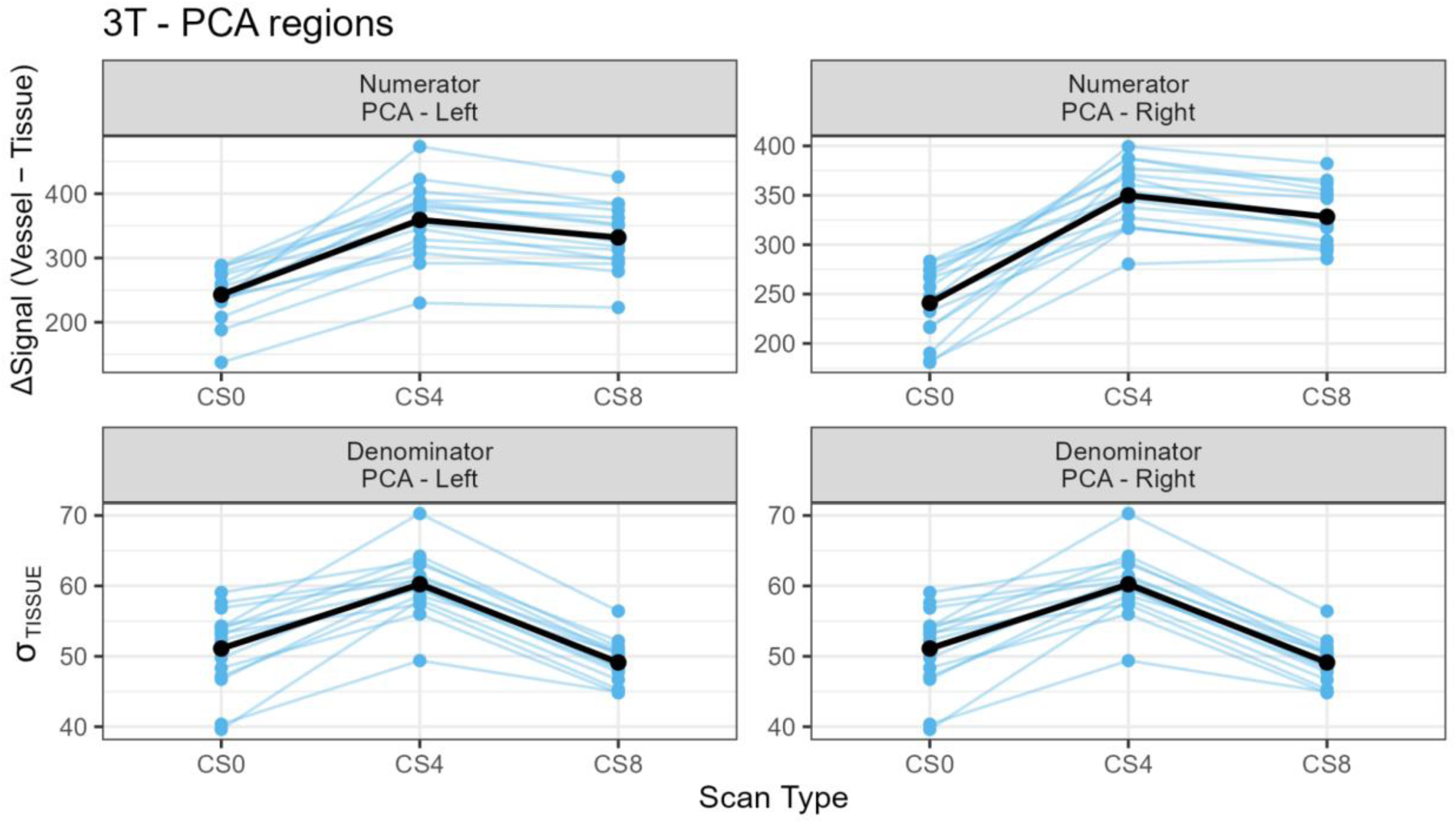
Comparison of acquisition types (CS0 and CS8) for select vessels at 3T separating the effect in numerator and denominator of the contrast-to-noise formula.

**Figure S5.**
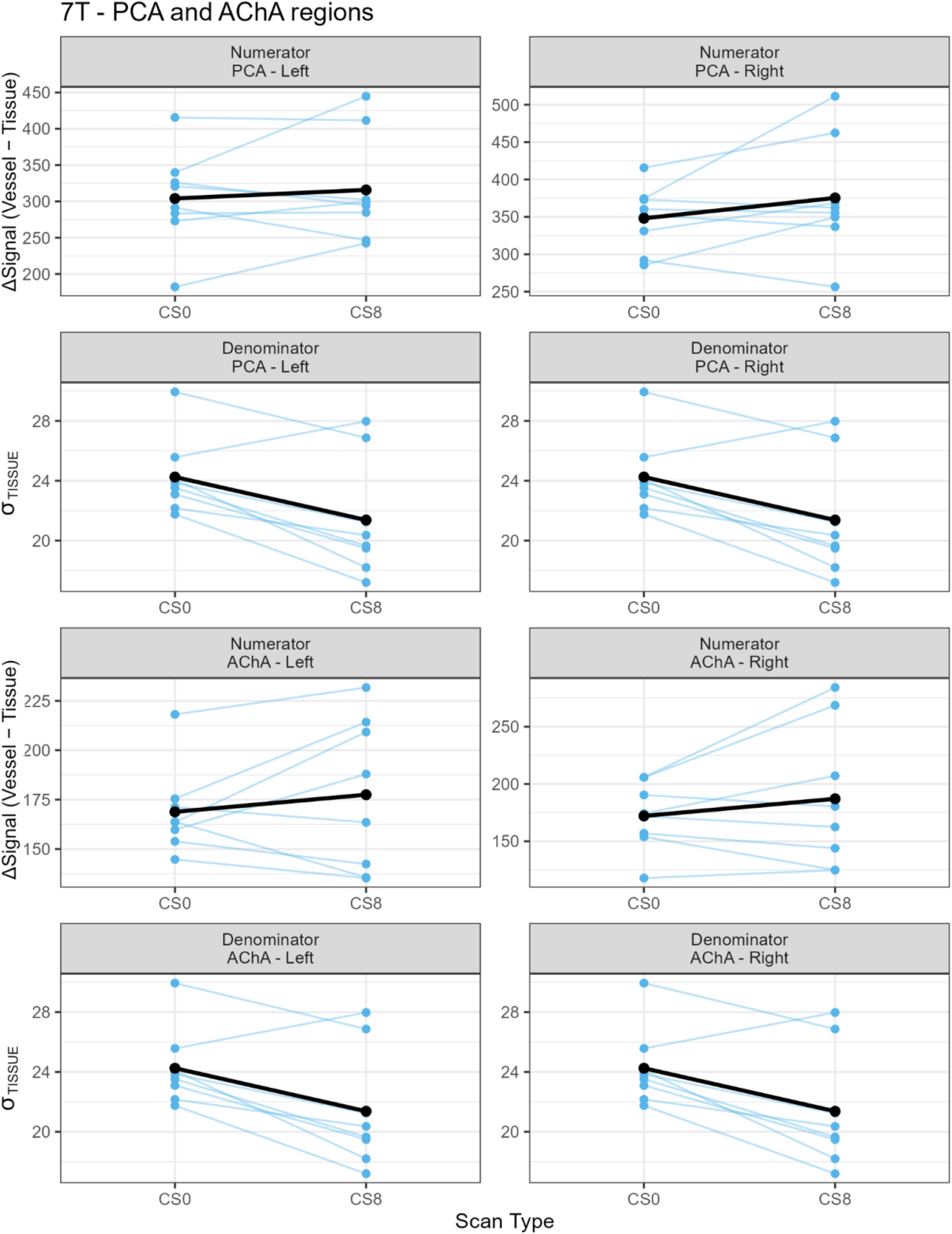
Comparison of acquisition types (CS0 and CS8) for select vessels at 7T separating the effect in numerator and denominator of the contrast-to-noise formula.

